# Excitoprotective effects of conditional tau reduction in excitatory neurons and in adulthood

**DOI:** 10.1101/2024.05.14.594246

**Authors:** Yuliya Voskobiynyk, Zhiyong Li, J. Nicholas Cochran, M. Natalie Davis, Nancy V.N. Carullo, Rose B. Creed, Susan C. Buckingham, Alicia M. Hall, Scott M. Wilson, Erik D. Roberson

## Abstract

Tau reduction is a promising therapeutic strategy for Alzheimer’s disease. In numerous models, tau reduction via genetic knockout is beneficial, at least in part due to protection against hyperexcitability and seizures, but the underlying mechanisms are unclear. Here we describe the generation and initial study of a new conditional Tau^flox^ model to address these mechanisms. Given the protective effects of tau reduction against hyperexcitability, we compared the effects of selective tau reduction in excitatory or inhibitory neurons. Tau reduction in excitatory neurons mimicked the protective effects of global tau reduction, while tau reduction in inhibitory neurons had the opposite effect and increased seizure susceptibility. Since most prior studies used knockout mice lacking tau throughout development, we crossed Tau^flox^ mice with inducible Cre mice and found beneficial effects of tau reduction in adulthood. Our findings support the effectiveness of tau reduction in adulthood and indicate that excitatory neurons may be a key site for its excitoprotective effects.

**SUMMARY:** A new conditional tau knockout model was generated to study the protective effects of tau reduction against hyperexcitability. Conditional tau reduction in excitatory, but not inhibitory, neurons was excitoprotective, and induced tau reduction in adulthood was excitoprotective without adverse effects.

## INTRODUCTION

The recent availability of amyloid immunotherapy for treatment of Alzheimer’s disease (AD) is a major advance. These treatments, however, only moderately slow the progression of disease (van Dyck et al., 2023; Sims et al., 2023) and will likely need to be combined with other drugs to achieve better efficacy. Drugs targeting microtubule-associated protein tau, the primary component of neurofibrillary tangles found in AD, are excellent candidates. Tau reduction using antisense oligonucleotides improved tau biomarkers in a Phase 1 clinical trial (Mummery et al., 2023; Edwards et al., 2023) and a Phase 2 trial is underway (NCT05399888). Thus, tau reduction is a promising therapeutic strategy for AD, and better understanding its effects is critical.

Tau reduction has strong beneficial effects in a variety of preclinical models, including mouse models of AD (Roberson et al., 2007; Ittner et al., 2010; Roberson et al., 2011; Nussbaum et al., 2012; Leroy et al., 2012; Lopes et al., 2016; DeVos et al., 2018). Similar beneficial effects were seen in rodent primary neuron and human induced neuron models of Aβ toxicity (Rapoport et al., 2002; Ng et al., 2024). Tau reduction appears to exert these benefits through resistance to hyperexcitability, which occurs in AD and AD models (Palop et al., 2007; Vossel et al., 2016; Lam et al., 2020). Tau reduction protects against epileptiform activity and other signs of hyperexcitability in AD mouse models (Roberson et al., 2007; Ittner et al., 2010; Roberson et al., 2011) and in both rodent and human neuron models (Voskobiynyk et al., 2020; Ng et al., 2024).

The protective effects of tau reduction against hyperexcitability extend beyond AD to models with seizures from diverse causes. Tau reduction protects against drug-induced seizures in wild-type mice (Roberson et al., 2007; Putra et al., 2020), and against seizures in models of stroke (Bi et al., 2017), Dravet syndrome (Gheyara et al., 2014), autism (Tai et al., 2020), and epilepsy due to potassium channel mutations (Holth et al., 2013). Tau reduction is also protective in *Drosophila* seizure models (Holth et al., 2013). Thus, tau reduction has broad excitoprotective effects, and therapies such as tau antisense oligonucleotides may have indications for a range of disorders associated with seizures (DeVos et al., 2013).

Many questions remain about the therapeutic potential of tau reduction. While tau reduction protects against hyperexcitability and seizures, better understanding its roles in excitatory vs. inhibitory neurons is critical for establishing the underlying mechanism. In addition, most studies of tau reduction to date have used traditional tau knockout models in which tau is reduced throughout development, which does not address whether tau reduction would be effective later in life after neurodevelopment. While most studies reported no adverse effects of tau reduction, a few have raised concerns about the safety of complete tau ablation (Lei et al., 2012; Ma et al., 2014), making it important to address the safety of tau reduction in adulthood.

To address these issues and enable future tau research, we generated a conditional tau knockout mouse model. Here, we use this model to compare effects of tau reduction in excitatory and inhibitory neurons and to study the effects of tau reduction in adulthood.

## RESULTS and DISCUSSION

### Generation of Tau^flox^ mice

To generate a model system with control over the location and timing of tau reduction, we developed Tau^flox^ mice for use with various transgenic Cre expression lines. Two *loxP* sequences were introduced through homologous recombination flanking exon 1 of *Mapt*, the tau gene (Fig. 1A, Fig. S1A–D), mimicking the design of the most widely used tau knockout line in which exon 1 is deleted, abolishing tau expression (Dawson et al., 2001). To confirm the expected function of the Tau^flox^ allele, a new tau knockout line was generated by breeding Tau^flox^ mice with CMV-Cre transgenic mice, which express Cre in all tissues, including germline. Resulting Tau KO mice had the expected loss of tau (Fig. S1E,F).

**Figure 1.**
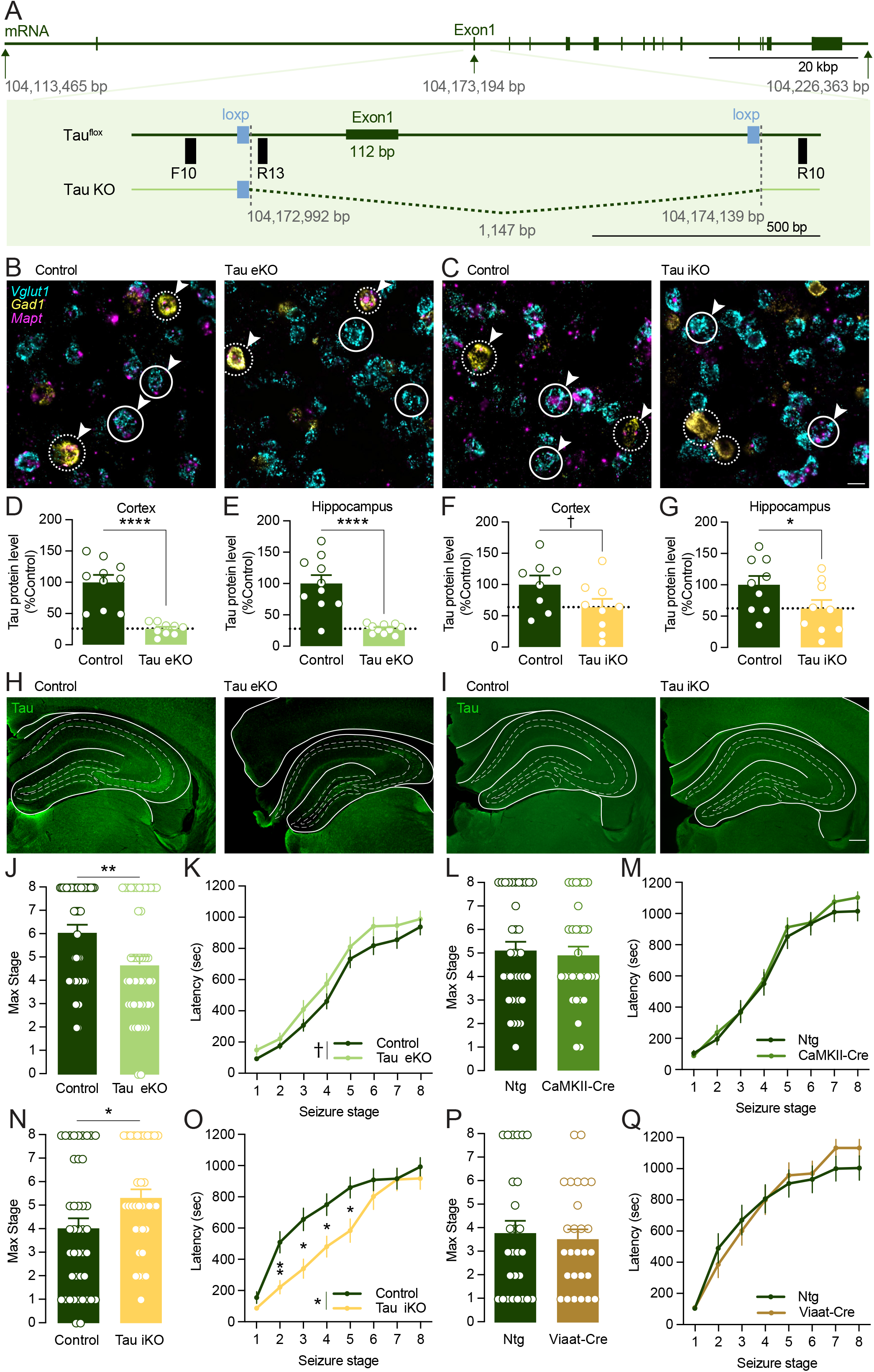
Differential effects of conditional tau reduction in excitatory vs. inhibitory neurons. (**A**) Schematic of the mouse *Mapt* gene (GRCm39 assembly) indicating the position of the *loxP* sites flanking exon 1 (the first coding exon) and the segment deleted by Cre recombinase. (**B, C**) Selectivity of tau reduction in Tau eKO (B) and Tau iKO (C) mice by single-molecule fluorescent in situ hybridization (smFISH) for excitatory neuron marker *Vglut1* (cyan, representative cells with solid circles), and inhibitory neuron marker *Gad1* (yellow, representative cells in dotted circles), and *Mapt* (magenta, in representative cells with arrowheads). **(D–G**) Degree of tau reduction in Tau eKO (D, E) and Tau iKO (F, G) mice, by western blotting in cortical (D, F) and hippocampal (E, G) homogenates (age 9–15 months, *n* = 9–10 per group, Student’s *t*-test **** P < 0.0001, ^†^P = 0.0861, *P = 0.0364. (**H, I**) Degree and localization of tau reduction in Tau eKO (H) and Tau iKO (I) mice by tau immunohistochemistry. (**J, K**) Lower seizure susceptibility in Tau eKO mice after PTZ, reflected by lower seizure severity (J, Mann-Whitney test, **P = 0.0044) and a trend toward longer latency (K, Two-way RM ANOVA, main effect of genotype ^†^P = 0.1218). Age 11–18m, *n* = 48–49 per group across 2 cohorts. (**L, M**) No effect of *CamKII*-Cre alone on seizure susceptibility after PTZ by seizure stage (L, Mann-Whitney test, P = 0.3832) or latency (M, Two-way RM ANOVA, main effect of genotype P = 0.5427). Age 9–15m, *n* = 37–31 per group. (**N, O**) Higher seizure susceptibility in Tau iKO mice after PTZ, reflected by higher seizure severity (N, Mann-Whitney test, *P = 0.0184) and shorter latency (O, Two-way RM ANOVA main effect of genotype *P = 0.0215, Holm-Sidak multiple comparisons test at stage 2 **P = 0.0054, stage 3 *P = 0.0107, stage 4 *P = 0.0407, stage 5 *P = 0.0407). Age 3–15m, *n* = 42–35 per group across 2 cohorts. (**P, Q**) No effect of *Viaat*-Cre alone on seizure susceptibility after PTZ by either seizure severity (P, Mann-Whitney test, P = 0.4670) or latency (Q, Two-way RM ANOVA, main effect of genotype P = 0.7983). Age 3–15m, *n* = 27 per group across 2 cohorts.

### Effects of tau reduction in excitatory vs. inhibitory neurons

Tau reduction has robust excitoprotective effects in various models with hyperexcitability, but the mechanisms remain uncertain. Establishing the relative roles of tau in excitatory and inhibitory neurons is a key step toward understanding these mechanisms. Not only are excitatory and inhibitory neurons likely to have fundamentally distinct contributions to hyperexcitability, but they are differentially affected by global tau reduction (Chang et al., 2021) and have differential susceptibility to tau pathology in AD (Fu et al., 2019).

To address this question, we crossed Tau^flox^ mice with cell type–specific Cre driver lines to make excitatory neuron knockout (Tau eKO) and inhibitory neuron knockout (Tau iKO) lines. We used CamKII-Cre mice, which express Cre selectively in excitatory forebrain neurons (Tsien et al., 1996), or Viaat-Cre mice, which express Cre selectively in inhibitory neurons (Chao et al., 2010). Tau eKO and Tau iKO mice had selective reduction of *Mapt* mRNA in either excitatory or inhibitory neurons, respectively (Fig. 1B,C). Total tau protein levels were reduced more substantially in the brains of Tau eKO mice than in Tau iKO mice, as expected given the greater number of excitatory neurons (Fig 1D–I). On behavioral screens, Tau eKO mice had a slight increase in exploratory locomotor activity (Fig. S2A), which has also been observed in global tauKO mice (Li et al., 2014), with no abnormalities on tests of anxiety-like behavior and working memory (Fig. S2B–D). Tau iKO mice, on the other hand, had normal locomotor activity and working memory (Fig. S2E,F) but increased anxiety-like behavior on elevated plus maze and light/dark box (Fig. S2G,H).

To test susceptibility to hyperexcitability in Tau eKO and Tau iKO mice, we challenged them with pentylenetetrazole (PTZ), a GABA_A_ receptor antagonist that induces stereotypical seizures that progress through well-defined stages. The maximal stage a mouse reaches and the latency to reach each stage serve as outcome measures of susceptibility to PTZ-induced seizures. Tau eKO mice reached a lower maximal seizure stage (Fig. 1J, Fig. S2I) and had longer latencies (Fig. 1K, Fig. S2J). We tested CamKII-Cre mice without a floxed allele to control for any tau-independent effects of Cre and observed no differences (Fig. 1L,M). These results indicate that selective tau reduction in excitatory forebrain neurons mimics the excitoprotective effects of global tau reduction, suggesting that these neurons are a key site for the effects of tau reduction.

We observed the opposite effect in Tau iKO mice, which showed increased susceptibility to seizures, with higher maximal seizure stage (Fig. 1N) and shorter latencies (Fig. 1O) after PTZ injection. These effects were not seen in Viaat-Cre controls without a floxed allele (Fig.1P,Q), indicating that they are tau-dependent. This increased seizure susceptibility is consistent with the increase in anxiety-like behavior (Fig. S2G,H), which are commonly observed together (Li et al., 2021; Qi et al., 2018). It is not surprising that manipulations in excitatory and inhibitory neurons have divergent effects on hyperexcitability, which has been observed with other genes (Deng et al., 2024). The observation of worsening hyperexcitability in Tau iKO mice indicates that inhibitory neurons are unlikely to be the primary site for the effects of tau reduction, although it does not rule out indirect effects on inhibitory neurons resulting from the effects of tau reduction in excitatory neurons.

### Inducible tau reduction in adult mice

Most studies showing benefits of tau reduction have used traditional global tau knockout lines in which tau is absent throughout development, but treatments for AD will require tau reduction in adulthood. To determine the effects of tau reduction in adulthood, we crossed Tau^flox^ mice with tamoxifen-inducible Cre transgenic (CreER™) mice. The resulting CreER™:Tau^flox/flox^ mice were injected with tamoxifen to produce tau adult knockout (Tau aKO) mice. In these mice, tau levels were reduced by ∼50% 2 weeks after tamoxifen injection and reached a nadir after ∼6 weeks (Fig. 2A). Tamoxifen reduced tau protein robustly in cortex and hippocampus by both western blot (Fig. 2B,C) and immunohistochemistry (Fig. 2D).

**Figure 2.**
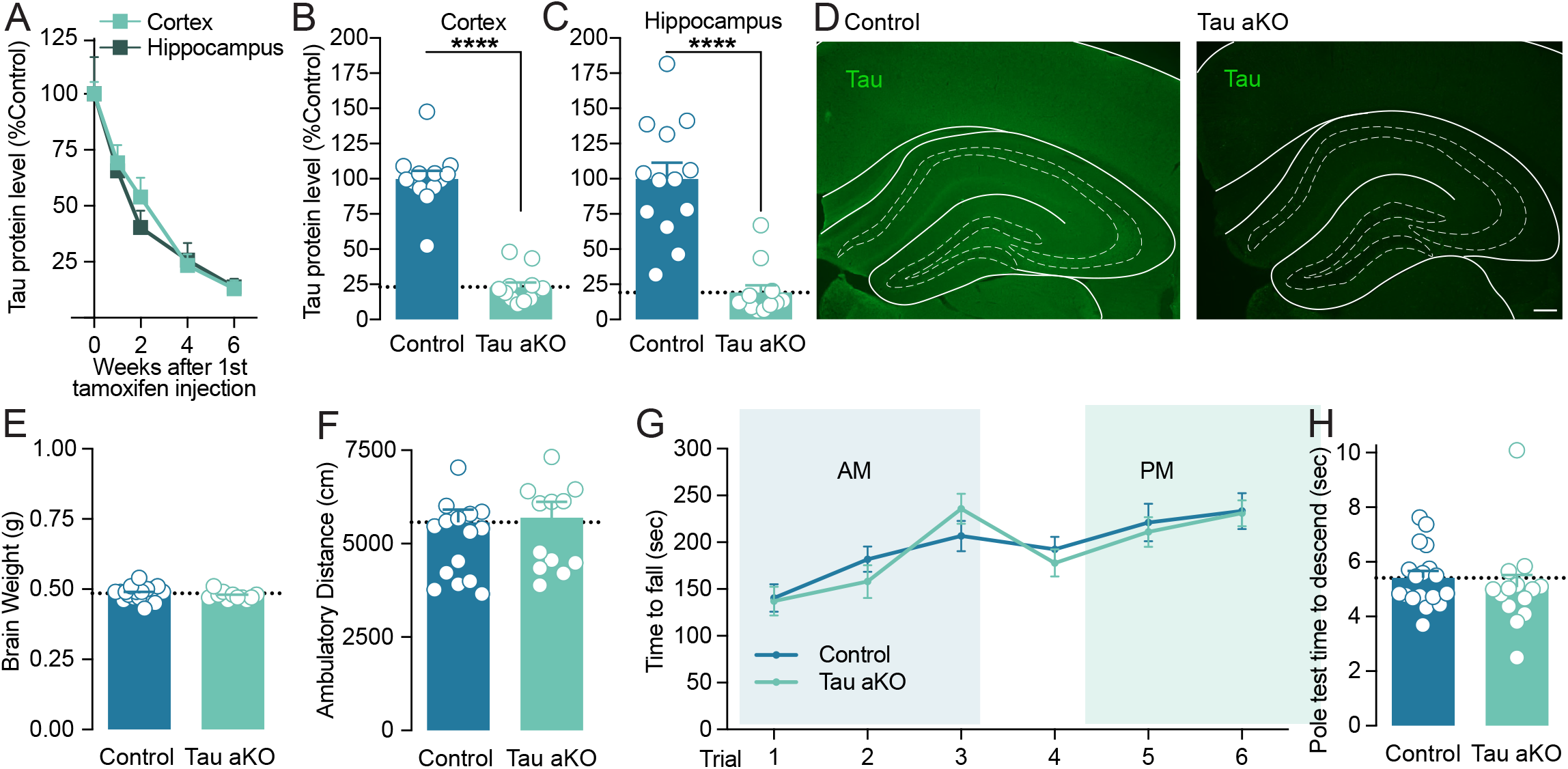
Conditional tau reduction in adulthood. Control (*Tau*^*flox/flox*^) and Tau aKO (*CreER*^*TM*^:*Tau*^*flox/flox*^) mice, age 2.5–4 months, received tamoxifen injections to activate CreER. **(A)** Time course of Tau reduction in Tau aKO mice. Mice were sacrificed at various time points after tamoxifen injection and Tau levels in the cortex and hippocampus were determined by Western blot. *n* = 3–5 mice per time point. **(B and C)** At steady state, 20 weeks after tamoxifen injection, mice were sacrificed, and Tau levels were determined by Western blot. Tau levels in aKO mice were 23% of control in the cortex (A) and 19% of control in the hippocampus (B). **** P < 0.0001 by Student *t*-test. **(D)** Tau immunohistochemistry 20 weeks after 1^st^ tamoxifen injection, showing robust inducible Tau reduction. **(E)** No difference in brain weight 20 weeks after tamoxifen (Student *t*-test, P = 0.3037). **(F)** No difference in ambulatory distance on open field testing 3 months after tamoxifen injection (Student’s *t*-test, P = 0.8163). **(G)** No difference in latency to fall from rotarod (Two-way RM ANOVA, main effect of Control vs. Tau aKO: P = 0.7718). **(H)** No difference in time to descend the pole test (Student’s *t*-test, P = 0.4981). Behavioral testing in F–H was 3 months after tamoxifen.

We next evaluated the safety of tau reduction in adult mice. Traditional Tau KO mice have slightly lower brain weights from an early age, likely due to a development effect (Li et al., 2014). There was no brain weight difference in Tau aKO mice (Fig. 1E), consistent with the idea that the smaller brain weight of conventional Tau KO mice is a developmental effect.

We also tested Tau aKO mice in behavioral assays. We focused on tests of motor function given reports of parkinsonism in some studies of aged conventional Tau KO mice (Lei et al., 2012). Tau aKO mice had no abnormalities in exploratory behavior or motor coordination in the open field (Fig. 1F), rotarod (Fig. 1G), or pole test (Fig. 1H).

In both cortex and hippocampus, Tau aKO mice had no changes in levels of various synaptic proteins, including Fyn, NR2B, Drebrin, PSD-95, and Synaptophysin (Fig. S3A,B). They also had no detectable upregulation of MAP1A or MAP2, nor any changes in tubulin (Fig. S3C,D). Acetylated tubulin, which can vary with changes in microtubule stability and was reported to be decreased in conventional Tau KO mice (Ma et al., 2014), was also unchanged in Tau aKO mice (Fig. S3C,D).

In summary, Tau aKO mice provide a model in which tau can be substantially reduced over the course of several weeks without notable adverse effects.

### Excitoprotective effects of tau reduction in adulthood

We next evaluated whether Tau aKO mice showed resistance to hyperexcitability, like traditional Tau KO mice. After challenge with PTZ, Tau aKO mice reached a less severe seizure stage (Fig. 3A) and had longer latency to reach seizure stages (Fig. 3B), indicating resistance to PTZ-induced seizures. To verify that these effects were not due to off-target aberrant activity of CreER™, we performed a similar experiment in CreER™ transgenic mice without the Tau^flox^ allele. CreER™ activation alone, without tau reduction, did not affect PTZ seizure susceptibility (Fig. 3C-D).

**Figure 3.**
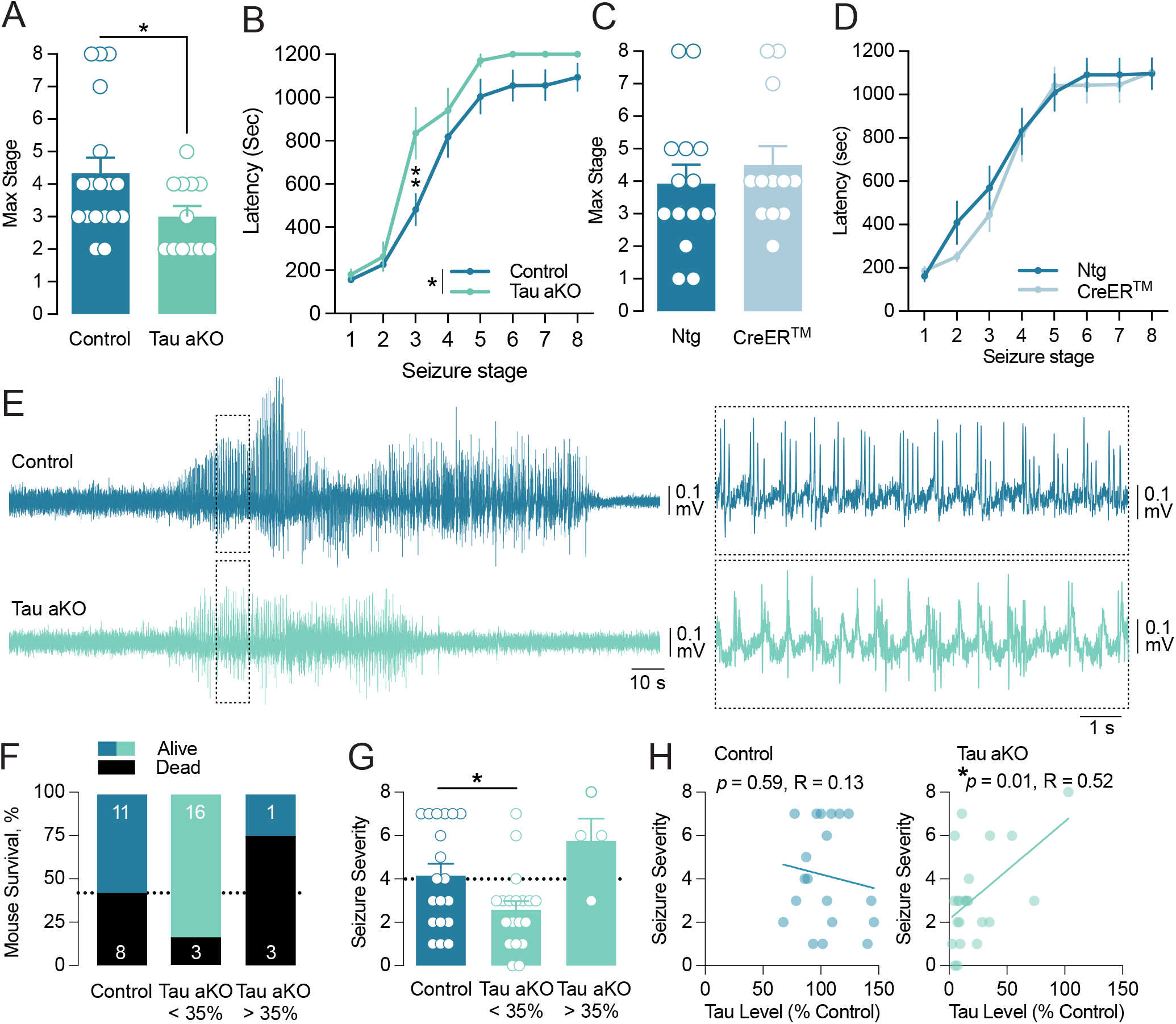
Tau reduction in adulthood increased resistance to PTZ- and kainic acid-induced seizures. **(A and B)** Control (*Tau*^*flox/flox*^) and Tau aKO (*CreER*^*TM*^:*Tau*^*flox/flox*^) mice, previously treated with tamoxifen, were intraperitoneally injected with 40 mg/kg PTZ. Tau aKO mice (A) reached a lower maximum stage of seizure severity (Student’s *t*-test, *P = 0.0482) and (B) had longer latency to reach seizure stages (RM two-way ANOVA, main effect of group *P = 0.0488, main effect of seizure stage ****P < 0.0001, Sidak’s multiple comparisons test **P = 0.0022 at seizure stage = 3). *n* = 12–18 mice per group. **(C and D)** Nontransgenic (Ntg) and CreER™ transgenic mice (both groups with wild-type, nonfloxed Tau), previously treated with tamoxifen, were intraperitoneally injected with 40 mg/kg PTZ. CreER™ activation had no effect on seizure resistance in the absence of tau reduction by either (C) maximum stage of seizure severity (Student’s *t*-test: P = 0.4959) or (D) latency to reach seizure stages (Two-way RM ANOVA, main effect of group P = 0.6258, main effect of seizure stage ****P < 0.0001). *n* = 12–14 mice per group. In A–D, mice were 2–5 months old when treated with tamoxifen, and PTZ testing was 4–5 months later. **(E–H)** Control (*Tau*^*flox/flox*^) and Tau aKO (*CreER*^*TM*^:*Tau*^*flox/flox*^) mice, previously treated with tamoxifen, were implanted with ECoG electrodes and intraperitoneally injected with 17.5 mg/kg kainic acid. (E) ECoG recordings of seizure activity from representative Control and Tau aKO mice after kainic acid injection. (F) Mortality in the first 48 h after kainic acid. (G) Seizure severity scores were lower than Controls in Tau aKO mice with effective tau reduction (ANOVA, P = 0.011; *P < 0.05 on posthoc testing). (H) Correlation between clinical score and residual Tau, showing that lower Tau levels are associated with lower seizure severity. In E–H, mice were 3–7 (average 4) months old when treated with tamoxifen, and kainic acid testing was 3–4 months later.

We next evaluated the resistance of Tau aKO mice to network hyperexcitability caused by kainic acid, a cyclic analog of L-glutamate and an agonist of ionotropic kainic receptors. Compared to PTZ, seizures induced by kainic acid are more variable and evolve over a longer period, so we used electrocorticographic (ECoG) recordings (Fig. 3E) and mortality as outcome measures. In this cohort, 4/23 Tau aKO mice did not show robust tau reduction (average tau levels 66% of control, vs. 14% of control in the other 19 mice), probably due to variation in tamoxifen injection. This variability in degree of tau reduction provided an opportunity to analyze the relationship between degree of tau reduction and resistance to kainic acid. In controls, 42% (8/19) of mice died in the first 48 hours after kainic acid. In Tau aKO mice that had effective tau reduction, mortality was only 16% (3/19), whereas in the Tau aKO mice with less tau reduction, mortality was 75% (3/4) (Fig. 3F). Using a scoring algorithm that also incorporated ECoG abnormalities, Tau aKO mice that had effective tau reduction had lower seizure severity than controls (Fig. 3G). We also analyzed the data without dichotomization, and in Tau aKO mice there was a strong correlation between the degree of tau reduction and the degree of resistance to kainic acid (Fig. 3H).

In summary, Tau aKO mice were resistant to both PTZ- and kainic acid–induced seizures, indicating an excitoprotective effect of tau reduction in adulthood.

### Conclusions and Perspectives

We describe here the generation of a new conditional tau knockout mouse model and its utility for studying the role of tau and the effects of tau reduction, a therapeutic strategy under investigation for AD. Our data support several conclusions about the mechanisms underlying previously published benefits of tau reduction. First, these benefits may arise from primary effects of tau reduction in excitatory neurons, where selective tau reduction mimicked the benefits of global tau reduction, and are unlikely to originate in inhibitory neurons, where selective tau reduction exacerbated seizure susceptibility (Fig. 1). Second, these benefits do not arise solely from developmental effects of the absence of tau, because we observed similar protection in Tau aKO mice in which tau is reduced in adulthood (Fig. 3). And third, these benefits do not require upregulation of MAP1A or MAP2, other microtubule-associated proteins, so are more likely due to a true loss of some function of tau (Fig. S3C,D). Tau^flox^ mice provide a tool that will be useful for answering diverse questions about tau function.

Given the excitation/inhibition imbalance that occurs in AD (Vico Varela et al., 2019), there is considerable interest in the roles of excitatory vs. inhibitory neurons, which are differentially affected by global tau reduction (Chang et al., 2021). We found that tau reduction in excitatory neurons was sufficient to reproduce the protective effects of global tau reduction (Fig. 1). This finding is consistent with that of another recent study that used a different Cre transgenic line (*Emx1*-Cre) to drive conditional tau reduction in excitatory neurons, showing excitoprotective effects in a model of the childhood epilepsy Dravet syndrome (Shao et al., 2022). Tau reduction in inhibitory neurons had the opposite effect, increasing seizure susceptibility (Fig. 1). The fact that global tau reduction (i.e. in both excitatory and inhibitory neurons) is excitoprotective indicates that the effects of tau reduction in excitatory cells are dominant over those in inhibitory neurons. Global tau reduction reduces action potential firing and excitation/inhibition ratio in cortical pyramidal cells (Chang et al., 2021), which is consistent with the reduced seizure susceptibility we observed in Tau eKO mice. Global tau reduction increases the excitability of inhibitory neurons, but this may be a secondary homeostatic response to reduced excitatory input (Chang et al., 2021), and further study is required to understand the effects of tau reduction in inhibitory neurons.

The safety of tau reduction has been a subject of considerable interest as it may seem surprising that reducing expression of an abundant brain protein would be well tolerated. Several studies have noted potential adverse effects of complete tau reduction in Tau KO mice, generally minor motor changes (Lei et al., 2012; Morris et al., 2013; Ma et al., 2014). However, several caveats should be noted. First, in many cases these findings have not been reproduced, and normal cognition has been repeatedly demonstrated in Tau KO mice (Morris et al., 2013; Li et al., 2014; van Hummel et al., 2016). Second, the described adverse effects of tau reduction are almost all limited to Tau KO mice which completely lack tau, whereas any potential tau reduction therapy would only partially reduce tau. Partial tau reduction, either in Tau^+/–^ mice (Roberson et al., 2007; Roberson et al., 2011), in mice treated with tau reducing therapies (DeVos et al., 2013; Wang et al., 2021), or in Tau aKO mice (Fig. 3), is sufficient to induce protective effects. Finally, studies with tau antisense oligonucleotides have moved questions about the safety of tau reduction beyond mouse models. Tau was reduced to ∼50% of normal levels without apparent adverse effects in nonhuman primates (DeVos et al., 2017). In a Phase 1 clinical trial with safety as the primary outcome measure, the highest doses reduced cerebrospinal fluid total tau by 51–56%. There were no associated serious or dose-limiting adverse effects, no discontinuations of drug dosing, and no abnormalities on safety magnetic resonance imaging (MRI) studies conducted after six months (Mummery et al., 2023). A phase 2 study in 735 participants (NCT05399888) launched in 2022 and continues, suggesting that the data safety monitoring committee has not identified major safety concerns to date.

This study has several limitations. First, neuronal subtypes are extensive, and while CamKII-Cre and Viaat-Cre are widely used tools to manipulate broad populations of excitatory or inhibitory neurons, additional Cre drivers could be explored in future studies to address these subtypes. Second, while our data suggest that a role for tau in excitatory neurons is likely dominant in mediating excitability effects, we did not explicitly test the role of glial tau. Finally, we focused in these initial studies on protective effects from seizures. Similar experiments could be conducted in the future focused on protective effects in AD mouse models. All of these are examples of potential future studies that are enabled by the availability of Tau^flox^ mice.

## METHODS AND MATERIALS

### Animals

All mice were housed in a pathogen-free barrier facility on a 12-hour light/dark cycle with *ad libitum* access to water and food (NIH-31 Open Formula Diet, #7917, Harlan). For postmortem analyses, mice were anesthetized by Fatal-plus (Vortech) and perfused with 0.9% saline. Mice were on a congenic C57BL/6J background. Mice were genotyped from tail tissue collected at weaning and verified again at death. Both male and female mice were used in experiments, and no sex-dependent effects were observed. Littermate siblings were used as controls in each experiment. Experiments were completed by blinded investigators in at least two separate cohorts, as indicated in figure legends. All experimental protocols were approved by the Institutional Animal Care and Use Committee of the University of Alabama at Birmingham.

### Generation of Tau^flox^ mice

*LoxP* sites were introduced into the mouse tau gene (*Mapt)* by homologous recombination. To generate the targeting vector, a 9kb fragment spanning exon 1 of *Mapt* was retrieved by homologous recombination into vector PL253, which contains the HSV-Thymidine kinase gene for negative selection. A *loxP* site was introduced 5’ of exon 1 along with an FRT-neomycin-FRT-*loxP* cassette 3’ of exon 1, plus a BamH I restriction site downstream of the 3’ *loxP* site to facilitate southern blot analysis (SFig 1A). After confirmation by sequencing, the targeting vector was linearized by restriction digests with Not I and gel purified.

The linearized targeting vector was transfected into C57BL/6-derived embryonic stem (ES) cells by electroporation at the University of Michigan Transgenic Animal Model Core. ES cells that were resistant to antibiotics G418 (indicating integration of neomycin cassette) and to ganciclovir (indicating absence of the HSV-TK cassette) were selected for further screening by Southern blot (SFig. 1B). Genomic DNA from ES cells was digested with restriction enzyme BamH I or Sac I (for 3’ and 5’, respectively) resolved on agarose gel, transferred under alkali conditions onto hybond XL membranes, and visualized using ^32^P-labeled probes. The probes were labeled with Prime-It primer labeling kit (Agilent) according to the manufacturer’s instructions. PCR was then used to confirm the presence of the 5’ *loxP* site and rule out the possibility of recombination between exon 1 of tau and 5’ frt site (SFig 1C). The following primers were used for PCR: F10, GCAATCACCTTCCCTCCATAACTAC; R10, ACCTTCGGAAGAGCAGTCGGGT; R13, TCAGCATCCCCACTAAAGCAGG.

ES cells with correct recombination were further examined for normal morphology and chromosome number and to rule out mycoplasma infection. Selected ES clones that contained the properly targeted (floxed) tau allele were microinjected into blastocysts at the UAB Transgenic & Genetically Engineered Model Systems Core Facility. Blastocysts were implanted in albino C57BL/6 mice to produce black-on-white chimeras.

Germline transmission of the floxed Tau allele was confirmed by PCR. Mice with the targeted allele were crossed with FLP transgenic mice (RRID:IMSR_JAX:009086) to delete the neomycin cassette, which was confirmed by PCR (SFig. 1D). This produced the Tau^flox^ mice used in this study. While this is the first full description of this line, these Tau^flox^ mice were shared with investigators for two prior reports (Narasimhan et al., 2020; Shao et al., 2022).

Tau^flox^ mice were crossed with germline-expressing CMV-Cre (RRID:IMSR_JAX:006054) mice to verify that Cre-mediated recombination deleted tau (SFig. 1E), producing Tau KO mice.

### Generation of excitatory or inhibitory neuron tau knockout (Tau eKO or Tau iKO) mice

Tau^flox/flox^ mice were crossed with CaMKII-Cre transgenic mice (RRID:IMSR_JAX:005359) for two generations to produce CaMKII-Cre:Tau^flox/flox^ (Tau eKO) mice and Tau^flox/flox^ controls. CaMKII-Cre drives expression in forebrain excitatory principal neurons (Tsien et al., 1996). Tau^flox/flox^ mice were crossed with Viaat-Cre transgenic mice (RRID:IMSR_JAX:017535) for two generations to produce Viaat-Cre:Tau^flox/flox^ (Tau iKO) mice and Tau^flox/flox^ controls. Viaat-Cre drives expression in GABAergic inhibitory neurons (Chao et al., 2010).

### Generation of adult tau knockout (Tau aKO) mice

Tau^flox/flox^ mice were crossed with CAGG-Cre-ER™ transgenic mice (RRID:IMSR_JAX:004682) (Hayashi and McMahon, 2002) for two generations to produce CreER™:Tau^flox/flox^ and Tau^flox/flox^ mice. All mice were injected intraperitoneally with tamoxifen (Sigma) at a dose of 160 mg/kg body weight. 100 mg tamoxifen powder was dissolved in 10 ml absolute ethanol with brief sonication, then 90 ml of corn oil (Sigma) was added, and the mixture was sonicated until it became completely clear. The tamoxifen solution was aliquoted and stored at −20°C.

### Western blot

Flash-frozen brain hemispheres were dissected into hippocampus and cortex, then homogenized in Tris-buffered saline with protease and phosphatase inhibitors. Protein concentrations were measured using BSA assay, and equal amounts of protein were loaded onto 4–14% Bis-Tris gel (Invitrogen) for electrophoresis, transferred onto PVDF-FL membrane (Millipore), immunoblotted with one of the following antibodies and visualized on Licor imager. Fyn (Cell Signaling rabbit polyclonal, catalog # 4023, RRID:AB_10698604); NR2B (Neuromab mouse monoclonal, catalog # N59/36, RRID:AB_2877296); Drebrin (Sigma rabbit polyclonal, catalog # D3816, RRID:AB_476905); PSD-95 (Cell Signaling rabbit monoclonal, catalog # 3409, RRID:AB_1264242); Synaptophysin (Millipore mouse monoclonal, catalog # MAB368, RRID:AB_94947); MAP1A (Abcam mouse monoclonal, catalog # ab11264, RRID:AB_2137390); MAP2 (Cell Signaling rabbit polyclonal, catalog # 4542, RRID:AB_10693782); Tubulin (Sigma mouse monoclonal, catalog # T5168, RRID:AB_477579); Acetylated tubulin (Sigma mouse monoclonal, catalog # T7451, RRID:AB_609894); Tau (DAKO rabbit polyclonal, catalog # A002401, RRID:AB_10013724).

### Immunohistochemistry

Saline-perfused brain hemispheres were fixed in 4% PFA for 24 hours, cryoprotected in 30% sucrose, and sectioned at 30 µm on a sliding microtome (Leica Biosystems). One set of serial sections was blocked with 10% normal goat serum and 10% milk for 1 hour at room temperature. The sections were incubated with anti-tau antibody (DAKO rabbit polyclonal catalog # A002401, RRID:AB_10013724) overnight at 4°C. The following day, sections were incubated with an Alexa Fluor-488-conjugated anti-rabbit antibody for 1 hour at room temperature. Sections were mounted on slides and coverslipped using ProLong Gold Antifade Mountant (Invitrogen). The immunostained sections were imaged at 4x using an epifluorescent microscope (Nikon ECLIPSE Ni).

### Behavioral testing

All behavioral tests were carried out at least one hour after the beginning of the light cycle. Mice were first brought into the testing room for acclimation at least 1 hour before experiments. Testing apparatuses were cleaned by 75% ethanol between experiments and disinfected by 2% chlorhexidine after experiments were finished each day. All mice were tested in all the behavioral tests in the same order. Investigators were blind to genotypes. Both male and female mice were used, and no sex-dependent effects were observed.

#### Open Field

Each mouse was placed in the corner of an open field (Med Associates) and allowed to walk freely for 15 minutes. Total and minute-by-minute ambulatory distance and rearing (vertical counts) of each mouse were determined using the manufacturer’s software. Average velocity of each mouse was manually calculated by dividing total ambulatory distance by total ambulatory time.

#### Rotarod

Mice were placed onto an accelerating rod (4–40 rpm in 5 min; Med Associates). The duration that each mouse remained on the rod was recorded. Each mouse was tested in two sessions on the same day with a 4-hour break between each session. During each session, each mouse was tested in three consecutive trials. The group average for each trial was reported.

#### Pole test

A wooden pole (3/8 inch diameter, 2 feet long) was placed perpendicular to the ground into a cage with 1-inch-deep bedding. Rubber bands were wrapped around the pole every 1.5 inches to provide grip. Each mouse was placed on the top of pole facing downwards and then released. All mice were trained for five trials on the first day and then tested with 5 consecutive trials the next day. The amount of time each mouse needed to walk down into the cage was recorded, and the best (shortest) trial test was reported.

#### Elevated Plus Maze

Elevated Plus Maze (Med Associates) has two open and two closed arms. Mice were placed in the hub of the maze and allowed to explore for five minutes. The time in each arm, as well as entrances to each arm, explorations, and head dips over the edge of the maze, were monitored by video tracking software (Med Associates). The maze was cleaned with 75% ethanol between mice and with 2% chlorohexidine at the end of each testing day.

#### Light/Dark Box

Light/dark testing was conducted in a chamber 20 cm L x 40.5 cm W x 22 cm H, split into one-third dark region (a black Plexiglass box with an opening) and two-thirds light region (exposed to room lights). Mice could freely explore for 10 minutes and were scored by a video tracking system (CleverSys).

#### Y maze

The Y-maze apparatus consisted of three 15-inch long, 3.5-inch wide, and 5-inch high arms made of white opaque plexiglass placed on a table. Each mouse was placed into the hub and allowed to freely explore for 6 minutes, with video recording. An entry was defined as the center of mouse body extending 2 inches into an arm, using tracking software (CleverSys). The chronological order of entries into respective arms was determined. Each time the mouse entered all three arms successively (e.g., A-B-C or A-C-B) was considered a set. Percent alternation was calculated by dividing the number of sets by the total number of entries minus two (since the first two entries cannot meet criteria for a set). Mice with 12 or fewer total entries were excluded from spontaneous alternation calculations due to insufficient sample size.

### PTZ-induced seizures

Pentylenetetrazole (PTZ, Sigma) was dissolved in PBS at a concentration of 4 mg/ml. PTZ at a dose of 40 mg/kg body weight was injected intraperitoneally. Immediately after PTZ injection, each mouse was placed into a cage for 20 minutes and observed by an investigator blind to its genotype, with video recording. The following scale was used: 0, normal behavior; 1, immobility; 2, generalized spasm, tremble, or twitch; 3, tail extension; 4, forelimb clonus; 5, generalized clonic activity; 6, bouncing or running seizures; 7, full tonic extension; 8, death (Racine, 1972; Löscher et al., 1991). The latency to reach each stage and the maximum stage each mouse reached were recorded.

### Kainic acid–induced seizures

EEG was used to assess electrographic seizures after kainic acid. Mice were anesthetized with 2.5% isofluorane. Six burr holes were drilled bilaterally through the skull, 2-, 4-, and 6-mm posterior to Bregma, and 2-, 4-, and 2-mm lateral to midline, respectively, using a dental drill equipped with a 1.0 mm drill bit. Three 1.6 mm stainless steel screws (Small Parts, Inc.) were screwed halfway into alternate holes. Then, an EEG electrode (Plastics One, Inc.) with 2 lead wires and a ground wire, cut to ∼1.5 mm, was inserted into the remaining three drill holes. The lead wires were placed bilaterally to record from each hemisphere. Dental acrylic was then applied to form a stable cap on the skull that cements the electrode. After the acrylic dried, the scalp was closed with skin glue (3M Vetbond). One week after electrode implantation, mice were transferred to specially constructed EEG monitoring cages where they were single housed. EEG data was amplified and digitized using Biopac Systems amplifiers (Biopac EEG100C) and AcqKnowlege 4.2 EEG Acquisition and Reader Software (BIOPAC Systems, Inc.). Data was stored and analyzed in digital format. Each cage was also equipped with an IR Digital Color CCD camera (Digimerge Technologies) that records each animal concurrently with EEG monitoring; recordings were acquired for review using security system hardware and software (L20WD800 Series, Lorex Technology, Inc.). All collected data was visually screened for epileptic events by an experienced observer blinded to genotype or treatment parameter. Abnormalities in the recordings indicative of epileptic activity were aligned chronologically with the corresponding video to confirm spiking or seizures.

Kainic acid (Tocris) was dissolved in PBS at a concentration of 2 mg/ml with intensive sonication. Kainic acid was injected intraperitoneally into each mouse at a dose of 17.5 mg/kg body weight. Immediately after injection, mice were returned to recording chamber and recorded for at least 24 hours. The following scale was used: 0, normal EEG; 1, EEG seizures ≤500 seconds/24hr; 2, EEG seizures 500–1000 seconds/24hr; 3, EEG seizures 1000–5000 seconds/24hr; 4, EEG seizures 5000–10,000 seconds/24hr; 5, EEG seizures ≥10,000 seconds/24hr; 6, death within 60–90 min; 7, death within 30–60 min; 8 death within ≤ 30 min.

### Fluorescent *in situ* hybridization (FISH)

FISH was performed using BaseScope and RNAscope Multiplex Fluorescent V2 Assays (Advanced Cell Diagnostics). Flash-frozen brain hemispheres were sectioned at 20 µm on a cryostat (Leica Biosystems), collected onto SuperFrost Plus slides (ThermoFisher Scientific), and immediately refrozen at −80°C. For pretreatment, tissue was removed from −80°C and immediately fixed with prechilled 10% neutral buffered formalin at 4°C, dehydrated with ethanol, and treated with hydrogen peroxide and protease III (ACD). Next, slides were incubated for 2 hrs at 40°C with a custom BaseScope probe (*BA-Mm-Mapt-2zz-st1-C1*, catalog #1322001-C1) targeting exon 1 of *Mapt* (130-215 of NM_001038609.3) to recognize the presence or absence of the floxed exon 1. Amplification of the probe was performed at 40°C, and development using BaseScope Fast RED A and B was performed at room temperature. Subsequently, RNAscope was performed by incubating tissue in a mixture of RNAscope probes (*Mm-Slc17a7-O2-C2*, catalog #501101-C2; *Mm-Gad1-C3*, catalog #400951-C3; ACD) for 2 hours at 40°C followed by fluorescent amplification. Slides were coverslipped with ProLong Gold Antifade Mountant with DAPI (ThermoFisher Scientific), and images were obtained with a Nikon AX confocal microscope.

### Data analysis

Data are presented as mean ± standard error of the mean. Unless otherwise specified, the experimental unit was each mouse. Sample sizes are included in each figure legend. No mice were excluded from the analyses. Statistical analyses were performed using Prism 10.0 (GraphPad).

## Acknowledgments

We thank James Black and Miriam Roberson for maintaining the mouse colony, genotyping, and tissue processing. We thank Dr. Andrew Arrant for training in behavior assessment and statistics and Dr. Travis Rush for assistance during PTZ experiments.

---

**Figure S1.**
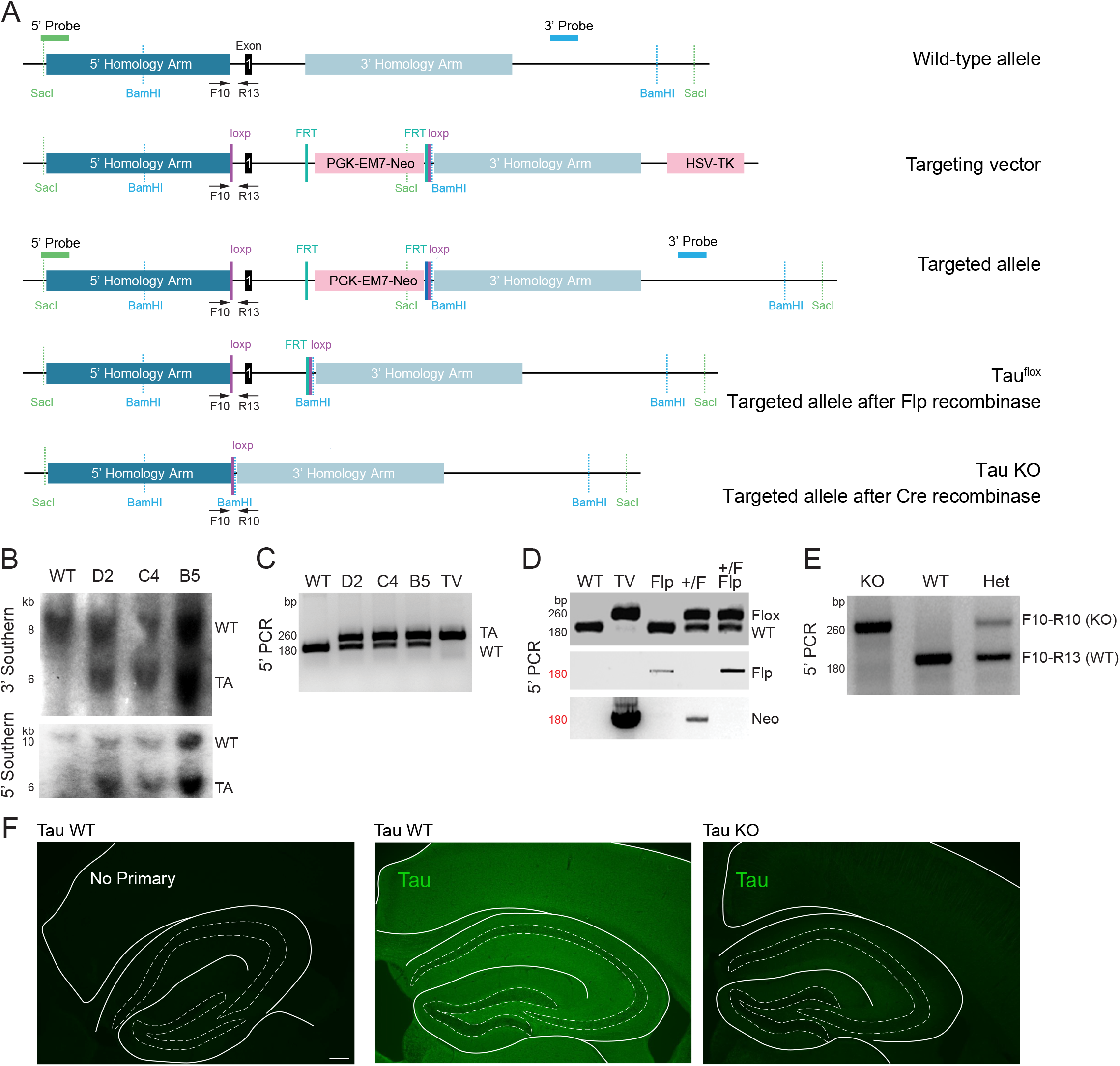
Generation of Tau^flox^ mice. **(A)** Schematic of wild-type Tau allele, targeting vector, targeted allele after homologous recombination, Tau^flox^ allele (targeted allele after Flp recombinase to remove Neo cassette), and Tau knockout (KO) allele (Tau^flox^ allele after Cre recombinase). **(B)** Southern blot confirmation of homologous recombination in three embryonic stem cell clones (D2, C4, and B5) after injection of the targeting vector. Top: southern blot with 3’ probe on genomic DNA digested with BamHI from WT ES cells and the three targeted clones. Bottom: southern blot with 5’ probe on genomic DNA digested with SacI. WT, wild-type allele; TA, targeted allele. TA bands run smaller due to insertion of novel restriction sites. **(C)** PCR genotyping with F10-R13 primers to confirm the presence of *loxP* sequence on 5’ side of Tau exon 1 in recombined ES cells. Lanes include DNA from WT ES cells, the three recombinant ES clones, and the targeting vector, TV. Top band represents the targeted allele (TA), which runs larger than the wild-type allele due to insertion of the *loxP* sequence. **(D)** PCR genotyping with F10-R13 primers (top), Flp primers (middle), and Neo primers (bottom) to confirm removal by Flp of the neomycin cassette from Tau^flox^ mice. WT, wild-type mouse tail DNA; TV, targeting vector. Flp, Flp transgenic mouse tail DNA; +/F, heterozygous Tau^flox^ mouse tail DNA. **(E)** PCR genotyping on tail DNA with F10, R10, and R13 primers to confirm deletion of Tau exon 1 by Cre recombinase. Lanes include wild-type (WT), Tau^+/–^ (Het), and Tau^−/–^ (KO) mice. **(F)** Tau immunohistochemistry of WT and KO brain sections showing expected loss of tau after Cre-mediated recombination.

**Figure S2.**
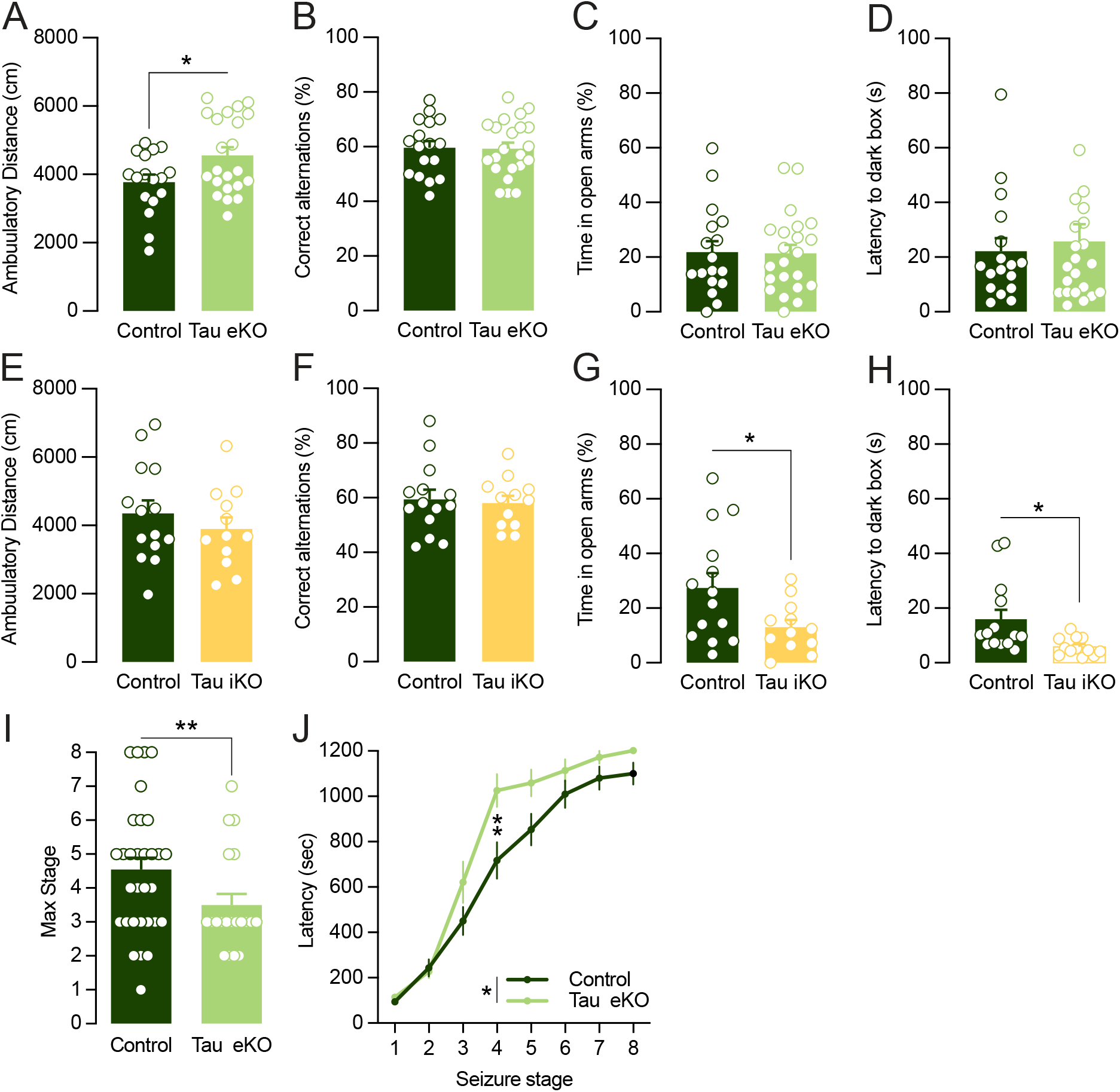
Behavioral effects of tau reduction in excitatory and inhibitory neurons and excitoprotective effects of tau reduction in excitatory neurons in aged mice. Behavioral screening of Tau eKO mice showing changes in (A) ambulatory distance in open field (Student’s *t*-test, *P = 0.0246) and no changes in (B) correct arm alternations in the Y maze, (C) time spent in open arms in elevated plus maze, nor (D) latency to the dark box in the light/dark box text (age 11–13m, *n* = 17–22 per group). Behavioral screening of Tau iKO mice showing no changes in (E) ambulatory distance in open field nor (F) correct arm alternations in the Y maze, but changes in (G) time in open arms in the elevated plus maze Student’s *t-*test, *P = 0.0343 and (H) latency to the dark box in the light/dark box test (Student’s *t*-test, *P = 0.0168; age 9–15m, *n* = 12–14 per group). (I, J) Lower seizure susceptibility in aged Tau eKO mice after PTZ (age 22–27m, *n* = 20–31 per group). (I) Lower maximum stage (Mann-Whitney test, *P = 0.0387). (J) Longer latency (Two-way RM ANOVA, main effect of seizure stage ****P < 0.0001, main effect of genotype *P = 0.0356, Holm-Sidak multiple comparisons test at stage 4 **P = 0.0014).

**Figure S3.**
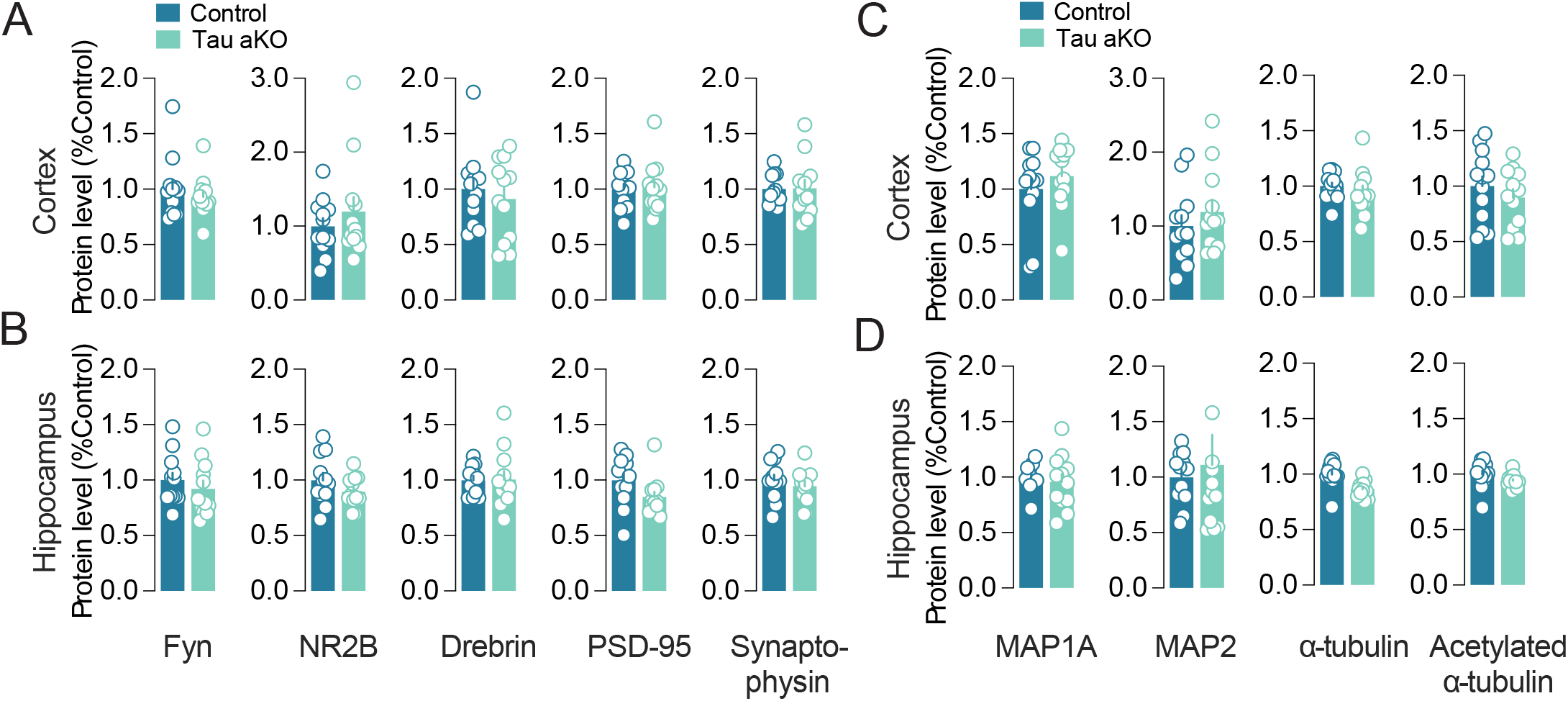
Tau reduction in adulthood does not alter synaptic, microtubule, or microtubule-associated proteins. Control (*Tau*^*flox/flox*^) and Tau aKO (*CreER*^*TM*^:*Tau*^*flox/flox*^) mice, age 2.5–4 months, received tamoxifen injections to activate CreER. **(A and B)** Several synaptic proteins were quantified by immunoblot in cortex (A) and hippocampus (B) of Tau aKO mice, showing no differences. **(C and D)** Several microtubule-related proteins were quantified by immunoblot in cortex (C) and hippocampus (D) of Tau aKO mice, showing no differences. Student’s *t*-test with Bonferroni adjustment for multiple comparisons.

